# Human fetal cerebellar cell atlas informs medulloblastoma origin and oncogenesis

**DOI:** 10.1101/2022.08.17.504304

**Authors:** Zaili Luo, Mingyang Xia, Wei Shi, Chuntao Zhao, Jiajia Wang, Dazhuan Xin, Xinran Dong, Yu Xiong, Feng Zhang, Kalen Berry, Sean Ogurek, Xuezhao Liu, Rohit Rao, Rui Xing, Lai Man Natalie Wu, Siying Cui, Lingli Xu, Yifeng Lin, Wenkun Ma, Shuaiwei Tian, Qi Xie, Li Zhang, Mei Xin, Xiaotao Wang, Feng Yue, Haizi Zheng, Yaping Liu, Charles B. Stevenson, Peter de Blank, John P. Perentesis, Richard J. Gilbertson, Jie Ma, Hao Li, Wenhao Zhou, Michael D. Taylor, Q. Richard Lu

## Abstract

Medulloblastoma (MB) is the most common malignant childhood brain tumor^1,2^, yet the origin of the most aggressive subgroup-3 form remains elusive, impeding development of effective targeted treatments. Previous analyses of mouse cerebella^3,4^ or human counterparts from frozen tissue nuclei^5^ have not fully defined the compositional heterogeneity of MBs. Here, we undertook an unprecedented single-cell profiling of freshly-isolated human fetal cerebella to establish a reference map delineating hierarchical cellular states in MBs. We identified a unique transitional cerebellar progenitor connecting neural stem cells to neuronal lineages in developing fetal cerebella. Intersectional analysis revealed that the transitional progenitors were enriched in aggressive MB subgroups, including group-3 and metastatic tumors. Single-cell multi-omics revealed underlying regulatory networks in the transitional progenitor populations, including transcriptional determinants HNRNPH1 and SOX11, which are correlated with clinical prognosis in group-3 MBs. Genomic and Hi-C profiling identified *de novo* long-range chromatin-loops juxtaposing HNRNPH1/SOX11-targeted super-enhancers to *cis*-regulatory elements of MYC, an oncogenic driver for group-3 MBs. Targeting the transitional progenitor regulators inhibited MYC expression and MYC-driven group-3 MB growth. Our integrated single-cell atlases of human fetal cerebella and MBs reveal potential cell populations predisposed to transformation and regulatory circuitries underlying tumor cell states and oncogenesis, highlighting hitherto unrecognized transitional progenitor intermediates predictive of disease prognosis and potential therapeutic vulnerabilities.

## Introduction

MB, a malignant pediatric brain tumor of the posterior fossa, is a highly heterogeneous tumor broadly comprised of Sonic Hedgehog (SHH), Group-3 (G3), Group-4 (G4), and WNT subgroups ^1,2^. Different MB subgroups can arise from diverse cell types or lineages in the developing cerebellum or brainstem and confer distinct treatment responses ^1–4,6^. The developing cerebellum is composed of distinct neural progenitor populations including progenitor cells in the cerebellar ventricular zone (VZ) that give rise to GABAergic neuronal lineage cells, and those in the rhombic lip (RL), which generate both granule cell progenitors (GCP) to form the external granule cell layer (EGL) and glutamatergic populations including unipolar brush cells (UBC) ^5^. The GCPs, UBC-lineage cells, and neural stem/progenitor cells are proposed as potential cells of origin for SHH-MB, G4-MB, and G3-MB, respectively^1–4,6^. The cells-of-origin of human G3-MB, the most aggressive subgroup associated with the worst prognosis and MYC activation^7,8^, have not been fully defined. Our understanding of MB tumor origins is largely informed by analyses of rodent models^3,4,9^. However, the human cerebellum has a 750-fold larger surface area than that of mouse with more primary progenitor populations^5,10^. Understanding the cellular heterogeneity across human cerebellar development is critical for decoding the developmental origin for G3-MB.

Here, we carried out single-cell transcriptomics profiling of whole cells from freshly isolated human fetal cerebella to define cellular hierarchy, transitional cell states, and their lineage trajectories during early cerebellar development. We compared these data to single-cell transcriptomes of MB subgroups to elucidate their developmental programs and identified a previously unrecognized transitional intermediate progenitor population in the fetal cerebellum as a potential cell of origin for aggressive MBs such as G3 tumors. Integrative single-cell multi-omics with 3D-genome architecture analyses further revealed unique tumor-driver networks and enhancer hijacking events correlated with *MYC* activation, pointing to potential therapeutic avenues.

## Results

### Human transitional cerebellar progenitors

We isolated fresh cerebellar tissues from aborted fetuses from post-conception weeks (PCW) 8 to 17 and profiled approximately 95,542 cells by single-cell RNA sequencing (scRNA-seq) after quality control and doublet removal^11^. Higher numbers of genes or read counts per cell were obtained in the scRNA-seq data than those from snRNA-seq experiments^5^ (Extended data Fig. 1a). Unsupervised clustering of individual cell transcriptomes visualized by t-SNE or UMAP^12^ identified 23 clusters (Fig. 1a, Extended data Fig. 1b, Supplementary Table 1). We annotated major cell types in human fetal cerebella by interrogating expression patterns of canonical markers for different cell lineages; the frequencies of cell types varied over time post-conception (Fig. 1a, Extended data Fig. 1c).

**Fig. 1.**
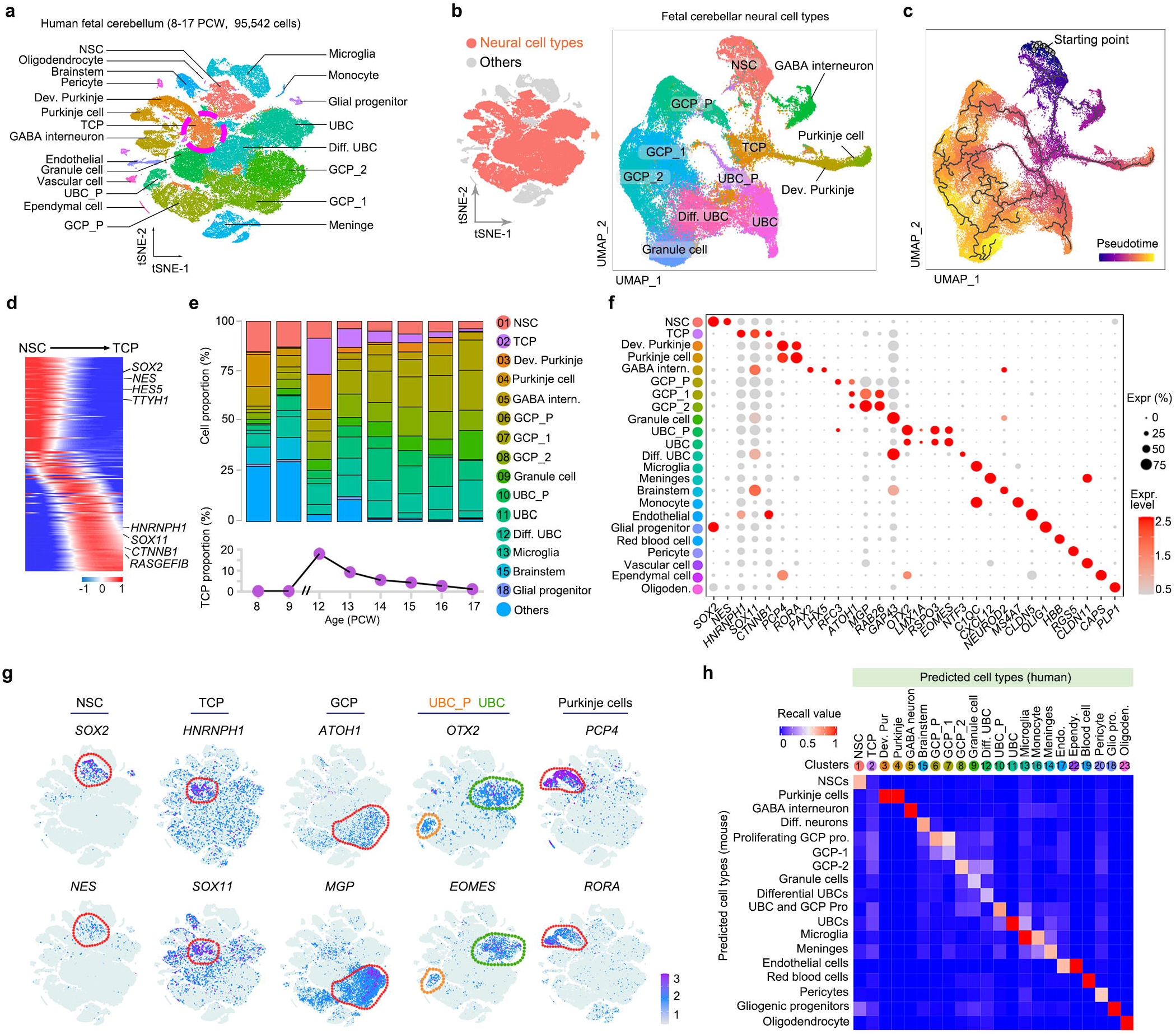
Single-cell atlas of early developing human fetal cerebella. **a**, t-SNE visualization of transcriptionally distinct cell populations from 95,542 single cells in PCW 8-17 human fetal cerebellar tissue. TCP are circled. **b**, Left: neural cell types (red) in t-SNE map of fetal cerebella. Right: UMAP visualization of neural cell types from fetal cerebella. **c**, Prediction of neural cell types fate transitions using Monocle 3. **d**, Heatmap with spline curves fitted to differentially expressed genes. Pseudo-temporal trajectories grouped by hierarchical clustering (k=3). **e**, Proportion of each cell population (upper) and the TCP proportion (lower) at indicated stages. **f**, Median-scaled ln-normalized expression of selected markers for identified fetal cerebellar cell states. **g**, Representative marker gene expression in the indicated cell types. **h**, A matrix for cross-validation with an SVM classifier of the identified cell types between the mouse and human fetal cerebella.

To focus on neural cell types, the presumed origin of MB, we investigated cell lineage trajectories using Monocle analysis^13^. We predicted the neural stem cells (NSC) population based on the stemness score as the start point and showed a trajectory through transitional cerebellar progenitors (TCPs) to the three main neuronal lineage branches, GCPs, UBCs, and Purkinje cells (Fig. 1b, c). STREAM^14^ confirmed this hierarchy (Extended data Fig. 1d). To define expression dynamics along the trajectory, we used Slingshot^15^ and found successional gene expression from NSCs to TCPs (Fig. 1d and Extended data Fig.1e). From PCW 8 to 17, there was an increase in the proportion of Purkinje, then GCP and UBC lineage cells (Fig. 1e), consistent with known dynamics^5^. TCP cells were abundant at PCW 12 and 13, but their frequency gradually decreased after PCW 14 (Fig. 1e). Sub-clustering indicated that a TCP population also expressed cell-cycle genes in G1/S and G2/M phases and proliferative marker Ki67 (Extended data Fig. 1f), suggesting that they are in a mitotic state. The TCP population was enriched in expression of *HNRNPH1* and *SOX11* and distinct from previously defined NSC, GCP, and UBC lineage cells (Fig. 1f, g and Extended data Fig. 1g), and the gene signature of TCP cells partially overlapped with that of RL cells^5^ (Extended data Fig. 2a-d). We further compared the populations we identified to published reference profiles of developing mouse cerebella^3^ using an algorithm for two-group classification^16^ and LIGER analysis^17^. Most human fetal cerebellar populations shared similarity with mouse counterparts; however, molecular features of the TCP population were not enriched in any known mouse cerebellar cell population (Fig. 1h and Extended data Fig. 2e-g).

### In vivo validation of TCP progenitors

We examined the expression of HNRNPH1 and SOX11, the most highly enriched TCP signature markers, in human fetal cerebella. HNRNPH1^+^ and SOX11^+^ TCP cells were increased in regions adjacent to the NSC (SOX2^+^) niche in the VZ from PCW 9 to PCW 12, a transitional period from the first to second trimester, but reduced progressively beginning at PCW 14 (Fig. 2a, b).

**Fig. 2.**
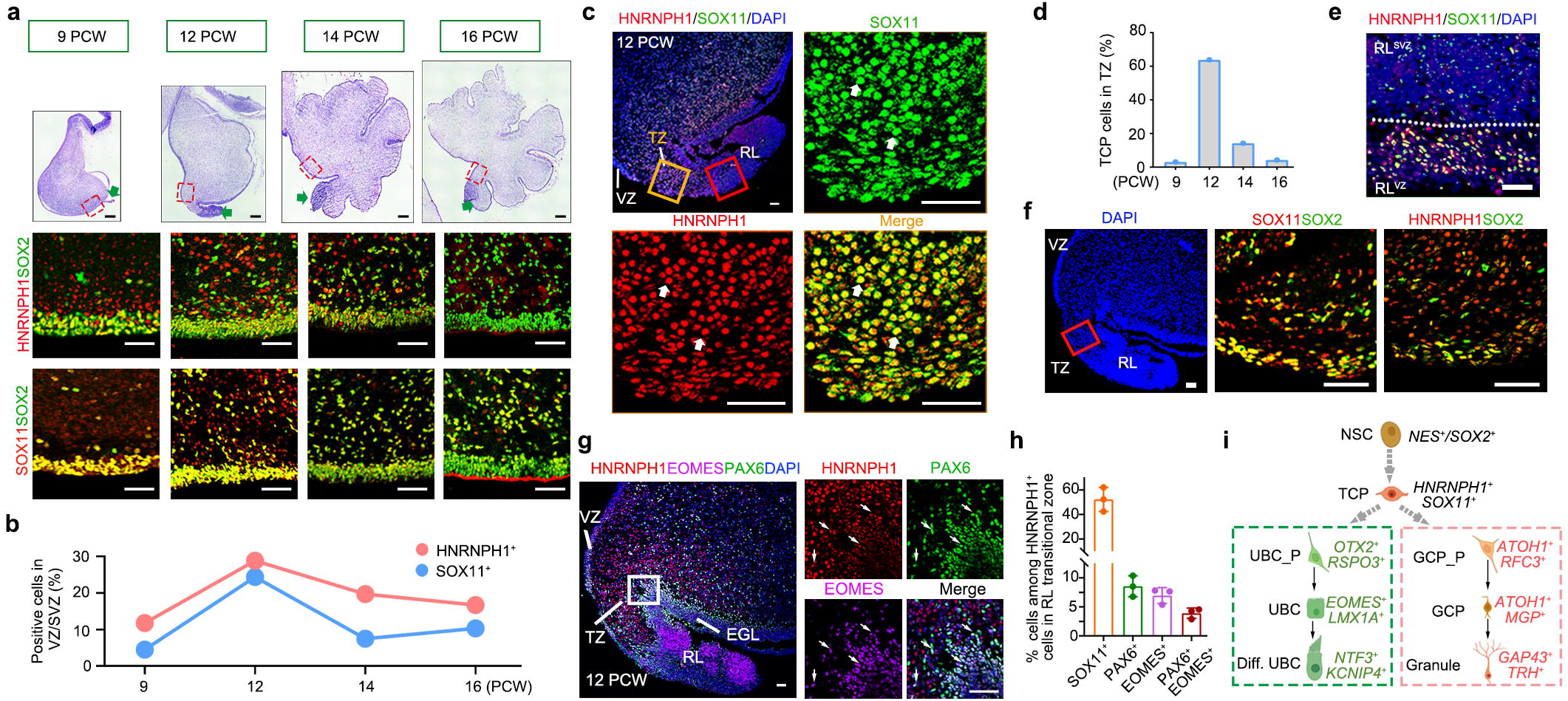
Identification of transitional progenitor intermediate in human fetal cerebella. **a**, Upper, H/E staining of midsagittal sections of PCW 9-16 human fetal cerebella. Arrows; the rhombic lip (RL). Middle and lower panels show SOX2/HNRNPH1 and SOX2/SOX11 immunostaining from the boxed areas in the upper panels at the corresponding stages, respectively. **b**, The percentage of HNRNPH1^+^ and SOX11^+^ cells at indicted stages (n= 2 biologically independent samples/stage). VZ, ventricular zone; SVZ, subventricular zone. **c**, Midsagittal sections of PCW 12 fetal cerebellum immunostaining for HNRNPH1 and SOX11 in the RL transitional zone (TZ) (yellow boxed). Arrows, co-labeled cells. DAPI (blue), counterstain. **d**, Proportion of TCP cells in the TZ at the indicated stages (n= 2 biologically independent samples/stage). **e**, Immunostaining for SOX11 and HNRNPH1 in the RL (RL^SVZ^ and RL^VZ^) from the red-boxed area in panel c at PCW 12. Scale bar: 100 μm. **f**, Midsagittal sections of PCW 12 fetal cerebellum immunostaining for SOX11 and HNRNPH1 in the TZ. DAPI (left), counterstain. Scale bar: 100 μm. **g, h**, Representative images (g) and the labeled cell proportions (h) in PCW 12 fetal cerebella stained for HNRNPH1, PAX6 and EOMES. Boxed areas are shown at a high magnification in the right panels. Arrows indicated triple-labeling cells. DAPI (blue), counterstain. Data are mean and SEM. **i**, Schematic model showing a potential transition through the TCP population into GCP and UBC lineages during the development of human fetal cerebella. Scale bars in **a**,**c**,**f**,**g**, 100 μm.

In the unique human fetal RL region ^5,10^, the TCP cell population with robust expression of HNRNPH1 and SOX11 was highly enriched in the RL transitional zone and RL^VZ^ region at PCW 12 compared with other stages, whereas low levels of TCP signature markers were detected in the RL^SVZ^ region (Fig. 2c-e). HNRNPH1 and SOX11 were also detected in a population of SOX2^+^ NSCs in the RL transitional zone (Fig. 2f).

Trajectory analysis predicted that TCPs may give rise to GCPs (ATOH1^+^) and UBCs (EOMES^+^) (Fig. 1c). Consistently, we detected a population of HNRNPH1^+^ cells that co-labeled with EOMES and PAX6 in the RL transitional zone (Fig. 2g,h), wherein PAX6^+^ progenitors can give rise to both GCPs and UBCs^18^. This suggests a potential lineage trajectory from TCPs to UBCs through PAX6^+^ intermediates. A population of ATOH1^+^ GCP progenitors in the posterior EGL layer were co-labeled with HNRNPH1 (Extended data Fig. 3a). Pseudo-temporal ordering of cell state evolution by Slingshot^15^ and STREAM^14^ revealed a trajectory initiated in NSC branches through the TCP subpopulation, which may serve as a precursor to generate GCP, UBC and Purkinje lineage cells (Fig2i and Extended data Fig. 3b,c).

### TCP-like cells in cerebellar MBs

To identify progenitor cells with molecular features of cerebellar MBs, we compared fetal cerebellar cell profiles to bulk transcriptomes of MB cohorts from the Children Brain Tumor Tissue Consortium using CIBERSORTx^19^. Consistent with previous observations^3,4^, the transcriptomic signatures of SHH MB from children and infants had strong similarity to GCP and child SHH tissues also exhibited weak similarity to TCP (Extended data Fig. 4a). The transcriptome profiles of G4 MBs resembled that of UBCs, whereas G3-MB cells (including MYC^high^ and MYC^low^ tumors) had the strongest similarity to human fetal TCPs, followed by UBC-lineage cells (Extended data Fig. 4a).

To further define the cellular identity of cerebellar MBs, we performed scRNA-seq and single-nuclei ATAC-seq (snATAC-seq) in matched tissues from 26 MBs (Supplementary Table 2). We also included in our analysis previously reported transcriptomics data^3^. TCP-like populations were identified in G3, G4, and SHH MBs as were tumor subtype-specific cell clusters (Fig. 3a). Unsupervised VECTOR trajectory analysis^20^ predicted that TCP-like cells were in an undifferentiated state (Extended data Fig. 4b-d). Reciprocal analysis of the overlaps between tumor cells and primary fetal cerebellar tissues using ProjecTILs^21^ revealed that MB tumor cells included TCP-like cell populations analogous to those in the human fetal cerebellum (Extended data Fig. 4-eh). TCP-like cells in different MB subgroups transcriptionally mimicked the normal TCP populations, whereas neoplastic cells in G3 and G4 MBs and in SHH MBs also had gene signatures similar to UBC lineage cells and GCPs, respectively (Fig. 3b). CIBERSORTx analysis^22^ showed that TCP-like populations were present in higher abundances in G3 MBs than in G4 and SHH MBs (Extended data Fig. 4i). TCP-like cells in MB tumors exhibited the TCP signatures (Fig. 3c) including expression of *HNRNPH1* and *SOX11* (elevated in a variety of cancers^23,24^) and *CTNNB1* (drives tumor formation in the cerebellum if normal lineage restriction is lifted^25^). TCP-like populations in G3, G4, and SHH MBs also expressed subgroup-specific signatures (Fig. 3c and Supplementary Table 3). These observations suggest that there is a potential tumorigenic evolution of TCP cells into specific neoplastic TCP-like cells in individual MB subgroups, possibly caused by distinct driver mutations.

**Fig. 3.**
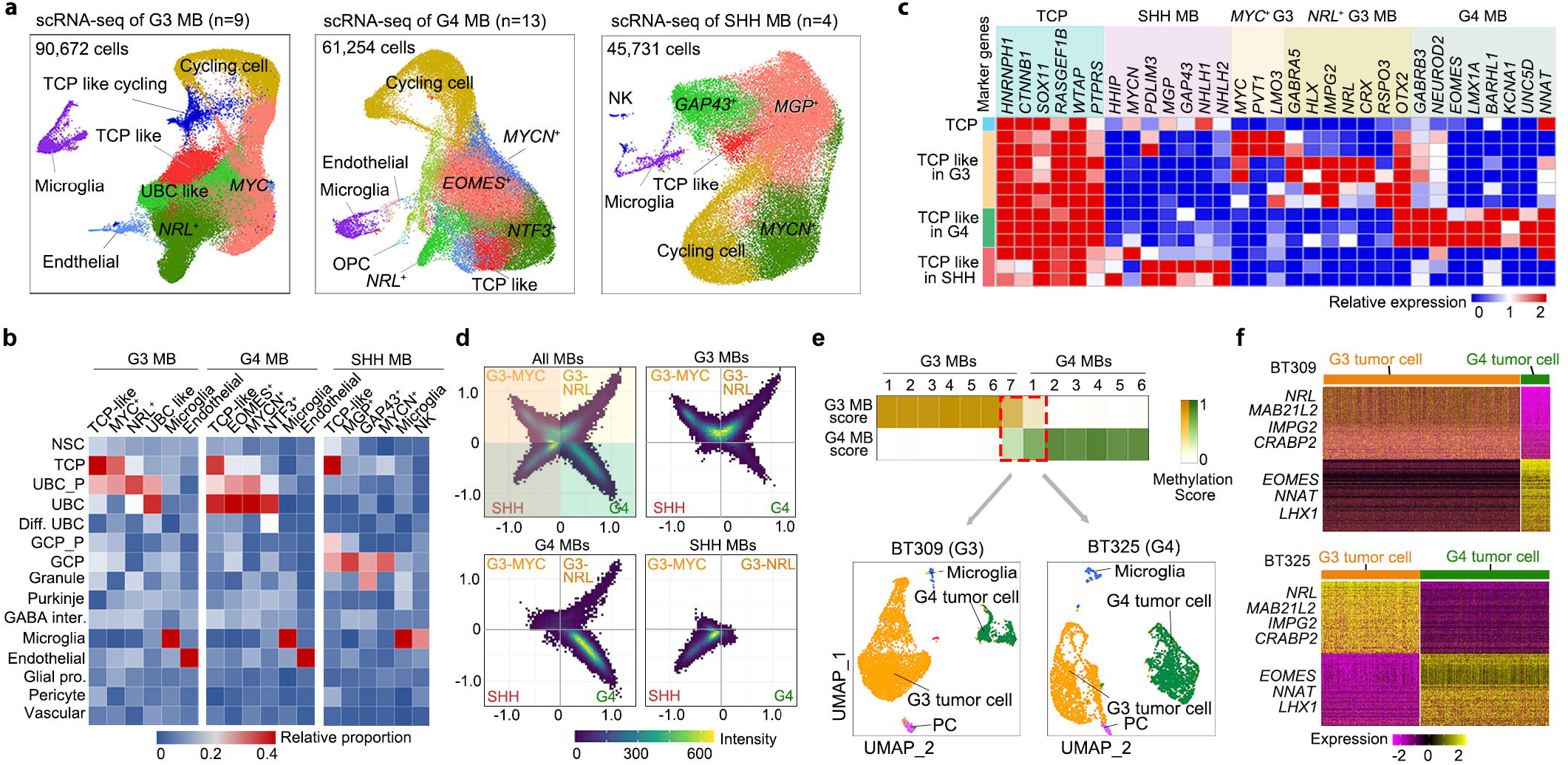
scRNA-seq reveals intermediate TCP-like progenitors in aggressive MBs. **a**, UMAP visualization of G3 (n= 9 tumors), G4 (n = 13 tumors), and SHH MB (n = 4 tumors). The cells are color-coded for indicated populations identified by representative marker genes. **b**, Heatmap of cell populations in human MB subgroups compared to normal human fetal cerebellar cell populations. **c**, Heatmap showing marker gene expression in pseudo-bulk normal fetal TCP and TCP-like tumor cells in SHH, G3, and G4 MBs. **d**, Cell-state plots of tumor subpopulations in indicated MB subgroups. The positions of cell states (dots) indicate relative meta-module scores, and color scales represent the count. **e**, Upper, DNA methylation-based subgroup prediction scores for G3 and G4 MBs. Samples G3#7 (BT-309) and G4#1 (BT-325) had a mixture of G3/G4 scores. Lower, UMAP distributions of G3 and G4 tumor cells from BT-309 and BT-325. PC, photoreceptor cells. **f**, Heatmap of the marker genes in BT-309 and BT-325 tumor cells showing mixed G3 and G4 cell populations.

Cell-state plots^26^ revealed specific enrichment of tumor cell states associated with MB subgroups (Fig. 3d). In contrast to the relatively restricted tumor cell state in SHH MB, we detected MYC^+^ and NRL^+^ cell states as well as a state with a G4 MB-like signature in G3 MBs (Fig. 3d). The G4 MB tumors were enriched in G4 MB-specific states and a cell state with an NRL^+^ G3 signature (Fig. 3d). These observations suggest that a cohort of G3 and G4 tumors have cells with either mixed or intermediate plastic states.

### Tumor cell populations in G3 and G4 MBs

G3 and G4 MBs share similar signature genes based on bulk transcriptome and DNA methylation profiles^4,27^. To evaluate the hypothesis that intermediate cell populations are shared between G3 and G4 tumors^4^, we analyzed two MBs (BT-309 and BT-325) that showed a mixture of G3 and G4 signatures based on a DNA methylation profiling (Fig. 3e). Unbiased single-cell clustering analysis of transcriptomes revealed distinct G3 and G4 tumor cell populations without substantial intermediate states expressing both G3 and G4 signatures (Fig. 3f and Extended Data Fig. 5a), and single-cell copy number variation (CNV) analysis confirmed that G3- and G4-like populations with distinct patterns were present within individual tumors (Extended data Fig. 5b). These data suggest that a set of G3 and G4 MB tumors might harbor a mixture of G3- and G4-specific cell populations rather than a G3/G4 cell state continuum.

### TCP frequency increases during metastasis

The BT-325 tumor, which harbored both G3- and G4- tumor cells, metastasized to the leptomeningeal surface of the brain. The metastatic tumor had increased frequencies of both TCP-like cells and MYC^+^ G3-like cells but a decrease in G4-like cells (Extended data Fig. 5c-e). There was enrichment in TCP-like and MYC^+^ G3-like states and gene signatures in the metastatic tumor coupled with a decrease in EOMES^+^ G4-like states when compared to the primary tumor (Extended data Fig. 5f,g). CNV analysis confirmed *MYC* gene amplification on chromosome 8 in the metastatic tumor in accordance with the higher level of *MYC* expression compared to the primary tumor (Extended data Fig. 5h,i). Similar observations were made in additional paired primary and metastatic G3 tumors (Extended data Fig. 5j-m) in concordance with the high rate of metastasis in G3 tumors^28^.

### Networks that drive TCP transformation

To decipher how dynamic accessibility at *cis*-regulatory elements (CREs) relates to the gene regulatory programs in TCP-like cells from aggressive MBs, we performed snATAC-seq of G3 and G4 MBs for which matched scRNA-seq data were available (Extended data Fig. 6a-c). By correlation of accessibility of promoter and gene body elements with target gene expression using ArchR^29^, we found that the positively correlated peak-to-gene pairs were mostly subcluster-specific (Extended data Fig. 6d-g).

To determine the temporal relationship between chromatin accessibility and gene expression, we ordered the peak-to-gene pairs based on their CRE accessibility as a function of pseudotime. In G3 tumors, the TCP-like cluster preceded the MYC^+^ cell cluster, which was followed by NRL^+^ cell cluster (Extended data Fig. 7a). Motif analysis indicated an enrichment of binding motifs for SOX11 and TWIST1 in the TCP-like cells, whereas TCF3 and MYC were enriched in the MYC^+^G3 cells, and NR2F1 and PAX5 motifs were enriched in the NRL^+^ G3 cells (Extended data Fig. 7a). Gene ontology analysis of G3-MB clusters showed an enrichment of epithelial development, epithelial-to-mesenchymal transition (EMT), and TGFb/BMP signaling in the TCP-like population (Extended data Fig. 7a). MYC+ populations by contrast were enriched in cell-cycle, cell migration, and Notch signaling, while NRL+ cells were enriched in photoreceptor cell development and Hippo signaling (Extended data Fig. 7a). The enrichment in EMT, TGFβ signaling, and cell migration likely contributes to the high metastatic potential of G3 MBs^30,31^.

In G4 MBs, the accessibility in TCP-like cells emerged prior to that of the KCNA1^+^ and EOMES^+^ cell clusters. An enrichment of epithelial cell development, neural progenitor cells, and UBC signatures was observed in TCP-like cells in G4 tumors (Extended data Fig. 7b). The KCNA1^+^ clusters were enriched in cell-cell adhesion, regulation of neuronal progenitors, and MAPK signaling, whereas the EOMES^+^ G4 subpopulation was enriched in neuronal development, HIF-1 and PI3K signaling (Extended data Fig. 7b). In EOMES^+^ G4 subpopulations, FOXG1 and LMX1A motifs were enriched, whereas RORA and PKNOX1/2 motifs were enriched in KCNA1^+^G4 subpopulations (Extended data Fig. 7b).

To identify positive transcriptional regulators that control gene expression in TCP-like-cell populations, we integrated snATAC-seq and scRNA-seq data to identify transcription factors with gene expression scores positively correlated with changes in accessibility of corresponding motifs^29^. TCP-like cell populations in G3 and G4 MBs were enriched in HLX, CRX, OTX2, BARHL1, and LMX1A in addition to TCP markers HNRNPH1 and SOX11 (Extended data Fig. 7c,d; Supplementary Table 3). CRE sites co-accessible with the promoters of potential drivers for MBs, including *OTX2* and *HLX,* were detected in TCP-like cells in G3 tumors, whereas *BARHL1* and *PAX6* were enriched in TCP-like cells in G4 tumors (Extended data Fig. 7e, f). Co-accessibility of regulatory CREs and target gene loci might contribute to expression of MB subtype-specific drivers and their oncogenic programs.

### Long-range enhancer hijacking in G3 MB

To identify the direct targets of SOX11 and HNRNPH1, we performed a Cut&Run genomic occupancy assay^32^ in patient-derived MYC-driven G3 MB tumor cell lines (MB-004 and MB-002) and non-transformed human NSCs and astrocytes (Fig. 4a). HNRNPH1 and SOX11 co-occupied enhancer and promoter regions near transcriptional start sites marked by activating histone marks H3K27ac and H3K4me3^33^ (Fig. 4b). H3K27ac signals were higher in G3 MB cells than NSCs or astrocytes (Extended data Fig. 8a and Supplementary Table 4). HNRNPH1 and SOX11 targeted common and unique genomic loci in G3 MB cells (Fig. 4a, boxed). Gene ontology analysis indicated that the unique targets in G3 MB cells were associated with the genes related to G3-MB oncogenesis, Ser/Thr kinase signaling, TGFβ/BMP signaling, and cell-cycle regulation (Fig. 4c). Expression of these genes was higher in G3 MB cells than control cells (Extended data Fig. 8b). Notably, HNRNPH1 and SOX11 targeted G3 signature genes *MYC* and *OTX2* in G3-MB cells but not in control cells (Fig. 4d). HOMER analysis of consensus sequence motifs associated with sites targeted by HNRNPH1/SOX11 revealed binding motifs for TGFβ/SMAD4, TCF3, and HIPPO/TEAD4 (Extended data Fig. 8c), which regulate tumor growth and metastasis^30,31,34^. Thus, our results suggest that HNRNPH1 and SOX11 might cooperate with these factors to regulate downstream gene expression that drives the G3 tumorigenic and metastatic programs.

**Fig. 4.**
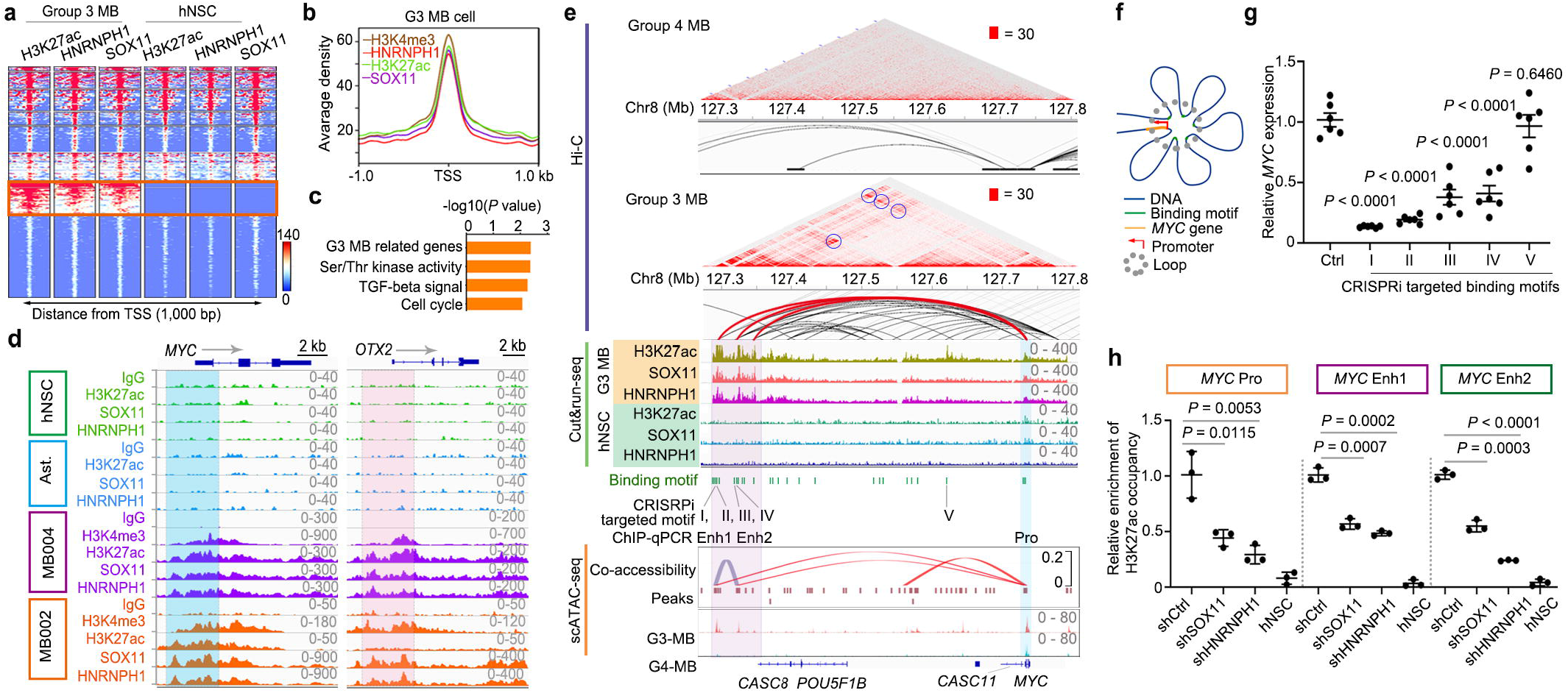
TCP regulators mediate long-range enhancer hijacking for G3-MB oncogenesis. **a**, Heatmaps of Cut&Run peaks with indicated antibodies in G3 MB cells and human NSCs showing ±1,000 bp around transcriptional start sites (TSS). **b**, Signals of H3K4me3, H3K27ac, HNRNPH1, and SOX11 peaks plotted relative to the TSS. **c**, Enrichment analysis of the genes co-occupied by HNRNPH1/SOX11 in G3 MB cells. *P*-value was calculated by one-sided Fisher’s exact test adjusted with multiple comparisons. **d**, Genomic tracks for H3K27ac, SOX11 and HNRNPH1 occupancy at *MYC* (left) and *OTX2* (right) loci of human NSCs and G3 MB cells (MB-004). **e**, Upper, Hi-C map of intrachromosomal interaction in G3-MB cells (MB-004) and G4-MB cells (UPN3550); middle, genomic occupancy; lower, pseudo-bulk genome co-accessibility in the *MYC* locus. Long-distance regulatory elements in human NSCs and G3 MB cells were highlighted. **f**, Proposed model for the contribution of enhancer hijacking to *MYC* expression. **g**, *MYC* expression after CRISPRi targeting SOX11/HNRNPH1-binding motifs (I, II, II, IV) and an internal motif V relative to control sgRNAs (n = 6 independent experiments/motif). **h**, Relative enrichment of H3K27ac occupancy on the indicated enhancers (Enh1 and Enh2) or promoter (Pro) in MB-004 cells transduced with non-targeting shRNA control (shCtrl), shSOX11, or shHNRNPH1 and in control hNSCs (n = 3 independent experiments). In g and h, data are shown as mean ±□SEM, unpaired two-tailed Student’s t-test.

To determine whether SOX11/HNRNPH1-occupied enhancers correspond to distal regulatory elements for activation of G3-MB driver genes, we performed Hi-C chromosome conformation capture in patient-derived G3 MB (MB-004) and G4 MB (UPN3550) cells and detected unique genomic looping in each line (Extended data Fig. 8d,e). We used NeoLoopFinder^35^ to reconstruct local Hi-C maps surrounding breakpoints. We identified unique structural variations and distinct sets of the interacting genomic loci involved in neo-loop formation at loop anchors in G3- and G4- MB cells (Extended data Fig. 8f,g). The neo-loop formation through inter-chromosomal translocation in G3-MB cells placed potential promoter/enhancer elements on a chromosome 11 segment close to the promoter of *PPP1R14A* on chromosome 19 (Extended data Fig. 8h), which has been shown to drive oncogenic RAS signaling in human cancers^36^. Expression of *PPP1R14A* was higher in the G3 MBs than SHH and G4 MBs (Extended data Fig. 8i). This suggest that inter-chromosomal translocation or structural variations might activate oncogenic drivers through enhancer-hijacking in G3 MBs.

Hi-C analysis indicated that the topologically associated domains in G3-MB cells harbored unique long-distance interactions with the enhancer and promoter regions of *MYC* (Fig. 4e). We also identified potential super-enhancers^37^ at the regulatory elements near the gene loci such as *OTX2, DUX4, CASC8* and *MYC* in G3-MB cells (Extended data Fig. 8j). By integrating the long-range interacting sites with enhancer occupancy of HNRNPH1/SOX11 in G3-MB cells and NSCs, we detected chromatin interaction loops linking the distal super-enhancer clusters bound by HNRNPH1/SOX11 upstream of *CASC8* to the active promoter/enhancer elements of the *MYC* locus in the G3-MB cells but not in NSCs (Fig. 4e). This suggests that long-range chromatin looping of super-enhancers juxtaposing to the *MYC* locus may promote *MYC* expression in G3-MB cells. CRISPR interference targeting of SOX11/HNRNPH1 binding motifs in the distal super enhancers, but not a control enhancer site, resulted in significant downregulation of *MYC* expression (Fig. 4f,g). In addition, ChIP-qPCR showed that the level of H3K27ac occupancies of enhancers and promoters of *MYC* or *OTX2* in G3-MB cells were reduced substantially when either SOX11 or HNRNPH1 was knocked-down (Fig. 4h and Extended data Fig. 8k). This long-range interaction loop was not detected in G4-MB cells (Fig. 4e), suggesting that a unique enhancer-hijacking event in the *MYC* locus drives oncogenesis in G3 MB.

### TCP-like signature genes in G3-MB growth

Based on the TCGA dataset, the level of TCP marker *HNRNPH1* was higher in G3 MBs than other MB subgroups, whereas *SOX11* expression was the highest in G4 MB (Extended data Fig. 9a). Notably, patients with high TCP scores (proportion of TCP)^19^ in G3 MB had worse prognosis than those with low TCP scores (Fig. 5a), whereas the prognostic impact was not observed in other MB subgroups (Extended data Fig. 9b). We detected HNRNPH1 and SOX11 in G3 and G4-MB tissues and co-expression of HNRNPH1 and SOX11 in a population of G3-MB cells (Fig. 5b). Proximity ligation assay showed that expression of HNRNPH1 and SOX11 was detected in the same foci in the nuclei of G3 tumor cells, and depletion of HNRNPH1 abrogated the signal (Fig. 5c). Consistent with their co-occupancy on CREs, these data suggest that HNRNPH1 may directly interact with SOX11 in G3-MB tumor cells.

**Fig. 5.**
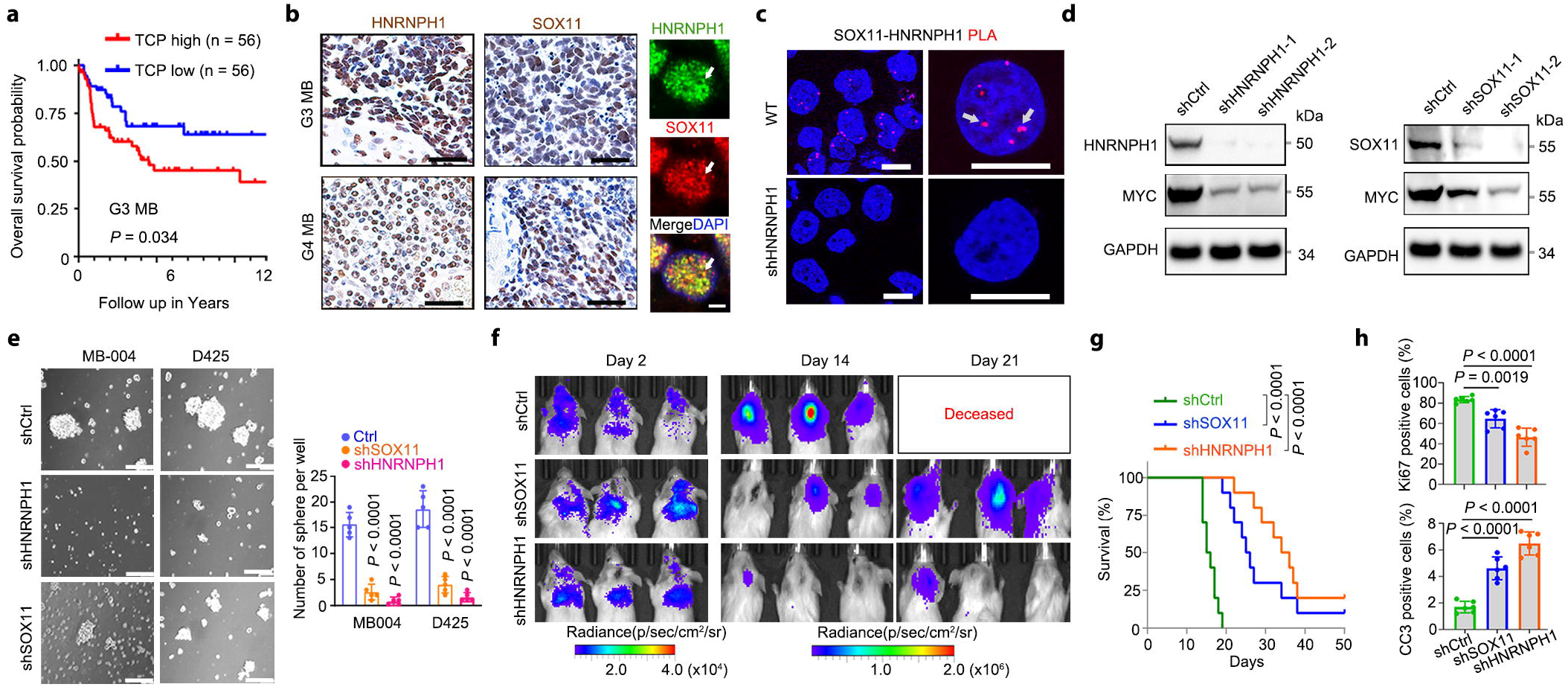
Targeting TCP regulators inhibits aggressive MB growth. **a**, Kaplan-Meier analysis of overall survival of patients with cerebellar MBs based on the TCP signature score in Cavalli’s MB cohorts. log-rank test. **b**, Left, representative images of human G3 (n = 12 tissue samples) and G4 (n = 9 samples) MBs stained for HNRNPH1 (left) and SOX11 (right). Scale bars: 100 μm. Right, representative images of HNRNPH1 and SOX11 co-immunostaining (arrows) in G3 MB (n = 3 samples). Scale bar: 2 μm. **c**, Representative images showed the proximity ligation assay of the HNRNPH1-SOX11 association (arrow) in MB-004 cells treated with shCtrl or sh HNRNPH1. n = 3 independent experiments. Scale bars: 10 μm. **d**, Representative immunoblots for MYC and HNRNPH1 in MB-004 cells (left) treated with control or shRNAs targeting *HNRNPH1* (left) or *SOX11* (right). n = 3 independent experiments. For gel source data, see supplementary Fig. 1. **e**, Images (left) and quantification (right) of tumor spheres formed by MB-004 or D425 cells treated with shCtrl, shSOX11, or shHNRNPH1. Scale bars: 200 μm. n = 5 independent experiments. **f**,**g**, Bioluminescent imaging (f) and survival curves (g) of NSG mice transplanted with MB-004 cells transduced with shCtrl, shSOX11, or shHNRNPH1 at indicated days (n=10 animals/group), log rank test. **h**, Quantification of tumor cell proliferation (upper, Ki67+) and apoptosis (lower, CC3+, cleaved caspase 3) in mice grafted with MB-004 cells transduced with shCtrl, shSOX11, or shHNRNPH1. n = 5 independent samples/treatment. In **e** and **h**, data are shown as the mean ±□SEM (bars) and individual score (dots); two-tailed unpaired Student’s *t*-test.

To determine the roles of HNRNPH1 and SOX11 in the growth of G3- and G4-MB tumor cells, we depleted tumor cells of HNRNPH1 or SOX11 using shRNAs. Silencing of *HNRNPH1* or *SOX11* in patient-derived G3-MB cells substantially reduced expression of MYC (Fig. 5d) and genes related to the G3-MB-associated signature while expression of neuronal differentiation genes increased (Extended data Fig. 9c-e). In addition, depletion of HNRNPH1 or SOX11 reduced tumor sphere formation in G3-derived MB-004 and D425 cells (Fig. 5e). Moreover, the growth of the patient-derived G3 MB-004 cells and of G4 MB UPN3550 cells was substantially reduced upon *HNRNPH1* or *SOX11* silencing (Extended data Fig. 9f), while depletion of either factor did not substantially impair the growth of NSCs or astrocytes. Silencing HNRNPH1 or SOX11 also significantly increased cell apoptosis (Extended data Fig. 9g). Furthermore, an *in vivo* xenograft analysis showed that SOX11 or HNRNPH1 depletion substantially inhibited the growth of G3-MB tumors and prolonged animal survival (Fig. 5f,g). The tumors depleted of HNRNPH1 or SOX11 had decreased proportions of Ki67^+^ proliferative cells and increased apoptosis (Fig. 5h, Extended data Fig. 9h). These results suggest that expression of the TCP signature genes *HNRNPH1* and *SOX11* is critical for the growth of aggressive G3-MB tumors.

## Discussion

In this study, we identified a previously unrecognized transitional progenitor population in the human fetal cerebellum. These cells were abundant during a narrow time window around the first-to-second-trimester transition stage, a period critical for neuronal lineage specification, proliferation, and migration^10,38,39^, and diminished thereafter. These progenitors had stem-like features of undifferentiated and transitory cell states with the potential to give rise to different cerebellar cell types including UBCs, GCPs, and Purkinje cell lineages. A recent study using snRNA-seq profiling of frozen fetal cerebella identified human RL cells but not the TCP subpopulation^5^. Our TCP gene signature partially overlaps that of RL cells^5^, yet the populations are distinct. Whole-cell scRNA-seq identifies cell types more representative of cell populations in the starting tissues than does snRNA-seq^40,41^, which might account for study differences.

Although SOX11^+^/HNRNPH1^+^ cells are sparsely distributed throughout mouse embryonic cerebella (Extended data Fig. 10a, b), they are not enriched in the RL region, in stark contrast to the enrichment of TCP cells in the human cerebellar RL transitional zone and RL^VZ^, the evolutionarily expanded region in humans^10^. This suggests that TCP cells represent cerebellar intermediate precursors similar to transit-amplifying progenitors involved in human neocortical expansion^42^. Such differences across species might explain why mouse model systems do not fully recapitulate human MB. By intersecting cellular states across developing fetal cerebella and MB subgroups, we discovered that TCP and tumor cell populations might be interconnected by tumor-subtype-specific transitory states. Our integrated single-cell omics and lineage trajectory analyses suggest that TCP-like cells might transition toward G3 tumors, serving as a potential cell of origin for G3 MB in a subset of tumors that form in infancy. Given the similarity between fetal and tumor cells does not necessarily indicate the tumor cell-of-origin, it is possible that malignant transformation occurs in other lineage precursors such as UBC lineage progenitors^43,44^ or through de-differentiation into TCP-like cells. The subgroup-specific transitory TCP-like progenitors within different MB subgroups may reflect intrinsic oncogenic mutations and cellular plasticity of TCP cells along distinct lineage trajectories, which may contribute to inter- and intra-tumoral heterogeneity as well as therapy resistance^45^.

Data from a recent study using SMART-seq or bulk RNA-seq implied a cell-state continuum among G3 and G4 MBs^4,46^. However, our unbiased single-cell clustering predicted that distinct populations of prototypical G3 and G4 tumor cells are present within G3/G4 tumors as opposed to a spectrum between G3 and G4 tumor cells. The limited gene sets for G3/G4 subtyping might diminish the distinction between cell types; a much higher number of cells and genes were assessed in our study. Our data indicate that G3 and G4 MB cell populations might not interconvert a subset of G3/G4 tumors, although our data does not exclude the possibility of an intermediate G3/G4 state.

Our single-cell profiling of paired primary and metastatic MBs revealed a substantial increase in the proportion of transitional TCP-cells and *MYC^+^* tumor cells in the metastatic tumors, suggesting that the TCP-like subpopulation and G3 tumor lineage cells drive metastatic tumor formation. We found that a TCP-like cell score is associated with poor prognosis in G3 MB but not in other subgroups. The higher proportions of the TCP-like cells in G3 MBs might contribute to the difference in survival outcomes. A TCP-like cluster was not detected in brainstem-derived WNT MB, which has the best prognosis of MB subgroups^47^ (Extended data Fig. 10c, d). Integrated scRNA-seq and snATAC-seq analyses identified transcriptional regulatory networks in TCP-like populations. Targeting TCP-like cells through depletion of HNRNPH1 or SOX11 inhibited the growth of the aggressive G3-tumor cells. Moreover, 3D-chromatin structure analysis revealed long-distance spatial looping of HNRNPH1/SOX11-bound superenhancers juxtaposed to *MYC* promoter/enhancer elements, which was uniquely present in *MYC*-driven G3-tumor cells but not G4 tumor or NSC cells. Thus, TCP cell identity determinants HNRNPH1 and SOX11, which are upregulated in multiple cancers^23,24^, not only define the TCP-like state but also hijack long-range superenhancers to promote expression of oncogenes. Together, our data provide insights into the potential origin, lineage plasticity, and human-specific nature of MB subtypes, as well as their intra- and inter-tumoral heterogeneity in malignancy and metastasis, while revealing a targetable vulnerability for therapeutic intervention of aggressive MB.

## Methods

### Human samples and tumor tissue collection

Human fetal and tumor tissues were obtained from the Children’s Hospital of Fudan University and XinHua Hospital at the Shanghai Jiao Tong University School of Medicine, and Obstetrics and Gynecology Hospital of Fudan University. Informed consents for the use of tissues for research were obtained in writing from donors or the patients’ parents in this study. The fetal and tumor tissue collections were approved by the individual institutional review board at the Children’s Hospital of Fudan University, XinHua Hospital at the Shanghai Jiao Tong University School of Medicine, and Obstetrics and Gynecology Hospital of Fudan University. Tumor tissue collections were approved by institutional review board at the Cincinnati Children’s Hospital Medical Center. Fresh cerebellar tissues from aborted fetuses and tumors after surgery were collected and digested by collagenase IV (2μg/ml, Thermo Fisher, #17104019) enzymatic dissociation for 20□min at 37□ after mechanically dissociation followed by single-cell profiling. Clinical information including age, sex, localization and MB subgroup are provided in Supplementary Table 1.

### Animal experiments

Immunodeficient NOD SCID gamma (NSG) 8-14 week old mice were obtained from the Cincinnati Children’s Hospital Medical Center (CCHMC) animal core. Mice of either sex were used and fed (4 or less mice per cage) in a vivarium with a 12 hrs light/ 12 hrs dark cycle. All studies complied with all relevant animal use guidelines and ethical regulations. The animal studies were approved by the IACUC (Institutional Animal Care and Use Committees) of the Cincinnati Children’s Hospital Medical Center, USA. In the xenograft model, MB tumor cells were transduced with lentivirus targeting HNRNPH1 or SOX11 for 20 hours, and 2×10^5^ cells suspended in 3μl PBS with 1μl Matrigel (Corning, #356234) were stereotactically injected into the cerebellum of NSG mice. Animals were monitored and bioluminescence images were captured weekly. Animals were removed from the study and were harvested when they exhibited >20% decrease in body weight based on the IACUC protocol. These limits were not exceeded in any of these experiments. Mice were housed at room temperature (20-23□) with a 12-h light-dark cycle set with lights on from 06:00 to 18:00 and with humidity between 30-80%. All the animal experiments were performed in accordance with the guidelines established by IACUC at the Cincinnati Children’s Hospital Medical Center. Animal survival endpoint is the date of the animal that died or was euthanized according to animal use guidelines. The limits of endpoints were not exceeded in any of the experiments.

### Medulloblastoma cell line culture

Medulloblastoma cell lines D425 (Cat# SCC290, Millipore sigma), DAOY (HTB-186, ATCC), D458 (CVCL_1161, Cellosaurus) and D283 (HTB-185, ATCC) were cultured in the DMEM/F12 (Thermo Fisher, #11320033) with 10% FBS and 2 mM L-glutamine and 1% penicillin/streptomycin. UPN3550 cells were isolated from a group 4 MB patient’s primary tumor tissue and cultured in the DMEM/F12 with 10% FBS and 2 mM L-glutamine and 1% penicillin/streptomycin, which was proved by the institutional reviewing board at the Cincinnati Children’s Hospital. MB004 and MB002 G3 MB lines^48^ were provided by Dr. Martine Roussel and cultured in neurobasal medium (Sigma; SCM003) with 2% B-27, 1 μg/ml heparin, 2 mM L-glutamine and 1% penicillin/streptomycin, 25 ng/ml FGF and 25 ng/ml EGF at 37°C in an atmosphere of 5% CO_2_.

### Immunostaining, immunohistochemistry and immunoblotting

The immunostaining procedures followed the method as previously described^49^. Briefly, cerebellar tissues were fixed with 4% PFA for 45 min and washed five times with PBS and dehydrated with 30% sucrose overnight, then blocked with OCT frozen embedding media (CRYO-4, Polarstat Inc.) and cryosectioned at 14 μm thickness. For adherent cells, cells were seeded on the coverslips and fixed with 4% PFA for 10 min and washed five times with PBS, then in blocking solution for 30 min. We used primary antibodies, including mouse anti-SOX2 (Santa Cruz Biotechnology; Cat#sc-365964), rabbit anti-SOX11 (Sigma, Cat#HPA000536; Thermo fisher, Cat#14-9773-82), rabbit anti-HNRNPH1 (Abcam, Cat#ab154894; Bethyl Laboratories, Cat#A300-511A), rat anti-EOMES (Invitrogen, Cat# 14-4875-52), mouse anti-PAX6 (Santa Cruz Biotechnology, Cat#sc-81649), mouse anti-ATOH1 (Thermo Fishe,; Cat#H00000474-M09), rabbit anti-c-Myc (Cell Signaling, Cat#5605), rabbit anti-Ki67 (Thermo Fisher, Cat#MA5-14520), rabbit anti-Cleaved Caspase 3 (Cell Signaling, Cat#9661) and mouse-anti BrdU (BD Bioscience, 1:500) antibody with proper dilutions. For BrdU staining, BrdU pulse-labeled (10 μM, 2 hours at 37□) cells were denatured with 0.1N HCl for 1 h in water bath at 37 °C. After denaturation, cells were neutralized with 0.1 M Borax, pH 8.5 (Sigma) for 10 min. Cells were washed with PBS 3 times and blocked with 5% normal donkey serum (Sigma-Aldrich) in wash buffer for 1 h at room temperature. Secondary antibodies conjugated to Cy2, Cy3 or Cy5 were from Jackson ImmunoResearch Laboratories. Tissues or cells were mounted with Fluoromount-G (SouthernBiotech) for microscopy. Immunofluorescence-labeled images were acquired using a Nikon C2^+^ confocal microscope. Cell images were quantified in a blinded manner.

For paraffin-embedded tissues, sections were dewaxed and hydrated using xylene and ethanol respectively. We performed antigen retrieval before permeabilization as previously described^49^. Slides were treated in 0.6% H2O2 in methanol for 30min at 37°C and blocked in 5% normal donkey serum in PBST for 1h at room temperature. SOX11 and HNRNPH1-expressing cells in MB tissues were quantified using the described methods ^50^. In brief, 0–5 denote different degrees (intensity and density) of IHC staining; 5 is the maximum and 0 is the minimum degree. The final score of the patients = SI (score of intensity) × SD (score of density).

For the western blot analysis, cells were lysed with RIPA lysis buffer (Millipore) supplemented with phosphatase and protease inhibitor cocktail (Roche). Protein concentration of each sample was determined by BCA assay using the BCA kit (Beyotime) according to manufacturer’s instructions and equal amounts (5-15 μl) were loaded and separated by 12% SDS-PAGE gel. PVDF membrane (Millipore) was used for gel transfer and the membrane was probed with primary antibodies as indicated, followed by secondary antibodies conjugated with HRP. The signal was detected with Super Signal West Pico/Femto Chemiluminescent Substrate (Thermo Scientific).

### Generation and processing of DNA methylation data

All single-cell MB patient samples sequenced in this study were analyzed using Illumina Infinium Methylation EPIC BeadChip arrays according to the manufacturer’s instructions. Data were generated from total genome DNA isolated from freshly frozen tissue samples. Medulloblastoma subgroup predictions were obtained from a web-platform for DNA methylation-based classification of central nervous system tumors (https://www.molecularneuropathology.org/mnp). Resulting assignment of samples to SHH, Group 3 and Group 4 subgroups were used for all downstream analyses. CNV analysis from EPIC methylation array data was performed using the conumee Bioconductor package (http://bioconductor.org/packages/conumee/).

### scRNA-seq and scATAC-seq using 10x Genomics platform

For scRNA-seq on the 10x Genomics platform, single cells were processed through the GemCode Single Cell Platform using the GemCode Gel Bead, Chip and Library Kits (10X Genomics) according to the manufacturer’s instructions. The concentration of the single-cell suspension was assessed with a Trypan blue count and the sample will be used if there are more than 90% viable cells. Approximately 10,000–30,000 cells per sample were loaded on the Chromium Controller and generated single-cell GEMs (gel beads in emulsion). GEM-reverse-transcription, DynaBeads clean-up, PCR amplification and SPRIselect beads clean-up were performed using Chromium Single Cell 3’ Gel Bead kit. Indexed single-cell libraries were generated using the Chromium Single Cell 3’ Library kit and the Chromium i7 Multiplex kit. Size, quality, concentration and purity of the cDNAs and the corresponding 10×library was evaluated by the Agilent 2100 Bioanalyzer system. Amplified cDNA and final libraries were assessed on an Agilent BioAnalyzer using a High Sensitivity DNA Kit (Agilent Technologies).

For snATAC-seq on the 10x Genomics platform, single-cell libraries were generated using the GemCode Single-cell instruments and the Single Cell ATAC Library & Gel Bead Kit and ChIP Kit from 10x Genomics, according to the manufacturer’s instructions. The samples were incubated at 37°C for 1 h with 10 μl of transposition mix (per reaction, 7 μl ATAC Buffer, and 3 μl ATAC Enzyme (10x Genomics)). Following the generation of nanoliter-scale GEMs, GEMs were reverse transcribed in a C1000 Touch Thermal Cycler (Bio-Rad) programmed at 72°C for 5 min, 98°C for 30 s, 12 cycles of 98°C for 10 s, 59°C for 30 s, and 72°C for 1 min, and held at 15°C. After reverse transcription, single-cell droplets were broken and the single-strand cDNA was isolated, cleaned up and amplified. Amplified cDNA and final libraries were assessed on an Agilent BioAnalyzer using a High Sensitivity DNA Kit (Agilent Technologies). All the libraries were sequenced on NovaSeq 6000 (Illumina) at a depth of approximately 400M reads per sample.

### scRNA-Seq processing and quality filtering

For 10X genomics datasets, we used Cellranger v5.0.1 to align reads to the human reference sequence. The raw base call (BCL) files were demultiplexed into FASTQ files. The FASTQ files were aligned to the reference human genome GRCh38 (hg38) to generate raw gene-barcode count matrices. When clustering multiple samples together, we aggregated the multiple runs together to normalize on sequencing depth and re-computed the gene-barcode matrices.

For quality control and normalization of scRNA-seq, we utilize the Seurat program (https://satijalab.org/seurat/articles/pbmc3k_tutorial.html) in R version 4.0.3 by reading in the data which are the reads in the output of the cellranger pipeline prom 10x, returning a unique molecular identified (UMI) count matrix. Low-quality cells were identified and removed from the datasets based on the cell with <200 genes expression and high mitochondrial gene content (5 s.d. above median). Doublets were detected and filtered using the R package DoubletFinder v.2.0.2 with default settings. The cells with low-abundance genes or genes expressed in <3 cells were also removed from the datasets. By defaulting in Seurat, we employ a global-scaling normalization method “LogNormalize” that normalizes the feature expression measurements for each cell by the total expression, multiplies this by a scale factor (10,000 by default), and log-transforms the result. Next, we apply a linear transformation (‘scaling’) that is a standard pre-processing step using all genes or variable genes. Then, we perform principal component analysis (PCA) to get the linear dimensional reduction after the data scaling.

### Clustering analysis, visualization and annotation

Clustering analysis was performed with the R package Seurat (v.4.0.3). Highly variable genes were detected using Seurat’s pipeline^51^, calculating average expression and dispersion for each gene, diving genes into bins and computing a z-score for dispersion within each bin. We used a z-score of 0.5 as the cut-off of dispersion, and a bottom cut-off of 0.0125 and a high cut-off of 3.0 for average expression. Linear dimensionality reduction was performed using PCA, and statistically significant principal components were selected using the elbow and jackstraw methods from Seurat. The clusters of cells were identified by a shared nearest neighbor (SNN)-modularity-optimization based clustering algorithm from Seurat. We then visualized these clusters using t-distributed stochastic neighbor embedding (t-SNE), uniform manifold approximation and projection (UMAP), or Monocle 3. Cluster cell identity was assigned by manual annotation using known cell-type marker genes and computed differentially expressed genes (DEGs) using the FindAllMarkers function in the Seurat package (one-tailed Wilcoxon rank sum test, P values adjusted for multiple testing using the Bonferroni correction. For selecting DEGs, all genes were probed provided they were expressed in at least 25% of cells in either of the two populations compared and the expression difference on a natural log scale was at least 0.2. Manual annotation was performed iteratively, which included validating proposed cell clusters with known markers and further investigating clusters for which the gene signatures indicated additional diversity.

### Pseudo-time cell state trajectory analysis

For the fetal cerebellum trajectory analysis, cells were grouped using ‘UMAP’ clustering algorithm. Cell state transition directions were inferred by Monocle 3, STREAM, or VECTOR programs which provides an unsupervised solution for determining the starting cells. For order_cells function in Monocle 3, the barcodes of selected clusters were normalized using Monocle dPFeature or Seurat to remove genes with low expression and perform PCA analysis on the remaining genes, for significant principal components (PCs) selection^52,53^. Differential gene expression analysis was performed using a generalized linear model, and the top 1,000 genes per cluster were selected and fit into a principal graph within each partition using the learn_graph function. For Slingshot cellular trajectory analysis ^15^ of fetal cerebella, the input matrix was filtered and normalized by the R package Seurat and cell types were annotated and provided as labels for Slingshot. For the single-cell pseudo-time trajectory in tumor tissues, cells were aggregated the multiple patients together to normalize on sequencing depth and re-computed the gene-barcode matrices using canonical correlation analysis (CCA).

### Deconvolution and overall patient survival analysis

CIBERSORTx^19^ was applied to perform the deconvolution analysis of the bulk and scRNA-seq tumor data against the human cerebellar clusters except mitotic cells. The transcriptomes of the tumor data (bulk RNA-seq or clusters of scRNA-seq) were used as the input mixture online (https://cibersortx.stanford.edu/runcibersortx.php), and the signature matrix input was the human fetal cerebellum cluster expression matrix after removal of cell cycle related genes (~1,400), ribosome biogenesis genes (~300), mitochondrial and apoptosis-related genes (~100), to avoid bias in the deconvolution process. Quantile normalization was disabled and 100–500 permutations for significance analysis were run.

Overall survival of human MB patients was right-censored at 12 years and analyzed by the Kaplan-Meier method. Patient cohorts were sub-grouped based on the TCP score (the estimated proportion of TCP). The TCP score was calculated based on the previously described scoring system with CIBERSORTx deconvolution analyses^19,54^ of the proportion of TCP cells against bulk transcriptomes of human MB subgroups (differentially expressed genes) from Cavalli’s MB cohort dataset. P values of survival curves were reported using the log-rank test.

### Cell cycle analysis of human scRNA-seq tumor samples

Cell cycle phase-specific annotations were used to define the cell cycle status for each individual cell^55^. We assign each cell a score using CellCycleScoring function in R version 4.0.5, based on its expression of G2/M and S phase markers. These marker sets should be anticorrelated in their expression levels, and cells expressing neither are likely not cycling and in G1 phase.

### Inferred CNV Analysis from scRNA-seq

Malignant cells were identified by inferring large-scale chromosomal copy-number variations (CNVs) in each single cell based on a moving averaged expression profiles across chromosomal intervals by inferCNV ^56^. We combined CNV classification with transcriptomic-based clustering and expression of non-malignant marker genes to identify malignant and non-malignant cells. Non-malignant cells displayed high expression of specific marker genes and no apparent CNVs.

### Filtering cells by TSS enrichment and unique fragments of the scATAC-seq

Enrichment of ATAC-seq accessibility at TSSs was used to quantify data quality without the need for a defined peak set. Calculating enrichment at TSSs was performed as previously described^57^, and TSS positions were acquired from the Bioconductor package from ‘TxDb. Hsapiens.UCSC.hg38.knownGene. Briefly, Tn5-corrected insertions were aggregated ±2,000 bp relative to each unique TSS genome-wide (TSS strand-corrected). The calculated TSS enrichment represents the maximum of the smoothed profile at the TSS. We then filtered all scATAC-seq profiles to keep those that had at least 1000 unique fragments and a TSS enrichment of 0.5. To minimize the contribution of potential doublets to our analysis, we removed snATAC-seq profiles that had more than 100,000 unique nuclear fragments.

### Gene regulatory network and motif enrichment analysis of scRNA-seq and scATAC-seq data

To characterize underlying gene regulatory network and infer transcription factor activities in our scRNA-seq dataset, we used the single-cell regulatory network inference and clustering package to identify gene regulatory modules and retain those with a cis-regulatory binding motif for upstream regulators. We merged scRNA-seq and scATAC-seq datasets to create a common peak set, and quantify this peak set in each experiment. We load the peak coordinates for each experiment and convert them to genomic ranges using the GenomicRanges::reduce function to create a common set of peaks to quantify in each dataset. We use the detailed settings and parameters as default according to Signac (https://satijalab.org/signac/). ArchR package^29^ was used for integrated scRNA-seq and scATAC-seq analyses according to default parameters, including quality control and cell filtering, dimension reduction, genome browser visualization, gene expression data preprocessing and cell annotation, DNA accessibility data processing, joint data visualization, differential accessibility and motif enrichment.

For nominating the marker genes and potential drivers, we utilized ArchR^29^ to identify the enriched TFs whose inferred gene scores are correlated to their chromVAR TF deviation z-scores. The gene scores were calculated based on the summed chromatin accessibility and normalized across all genes to a user-defined constant (default of 10,000) according to the ArchR package ^29^.

Based on the gene scores and positive TF-regulators identified from ArchR, we nominated top 30 TFs or highly expressed genes (excluding non-coding genes or ribosomal proteins) as potential drivers or marker genes.

### Two-Dimensional Representation of SC-Derived Cell States and Group 3/4-B/C score

Tumor cell clusters were used for computing subtype expression scores for each tumor cell in the datasets as previously described ^26^. Cell clusters were separated into G3-MYC, G3-NRL, G4- and SHH-clusters. To visualize the enrichment of subsets of cells, across the two-dimensional representation, we calculated for each cell the fraction of cells that belong to the respective subset among its 100 nearest neighbors, as defined by Euclidean distance, and these fractions were displayed by colors. In addition, we used the previously defined group 3/4 B/C score system based on selected G3- and G4-expressing genes (top 30 genes from each metaprogram) ^4^ for the overlapping cell-state analysis in G3 and G4 MBs.

### Generation of Hi-C libraries and analysis

MB004 (G3 MB) and UPN3550 (G4 MB) cells were processed for Hi-C library construction using the Arima Hi-C Kit following the manual (Arima Genomics, # A510008). Briefly, five million cells were cross-linked with 1% formaldehyde for 10 mins at room temperature and then quenched with 0.2M Glycine. Cell pellets were washed with cold PBS and lysed with lysis buffer to release nuclei and then permeabilized and in situ digested. KAPA Hyper Prep kit was used for library amplification (KAPA, KK2802). Hi-C libraries were sequenced 2×150bp on a NovaSeq 6000 instrument (Illumina). Juicer were used to process raw reads and generate Hi-C contact matrices (.hic files), aligning to reference genome hg38 to generate Hi-C contact matrices (.hic files). Contact matrices were visualized using Juicebox.

Bam files were used as input, with low-quality reads filtered out. We used Peakachu and diffPeakachu to call and compare loops in Hi-C data from both G3 MB and G4 MB cell lines in Hi-C data, then used diffPeakachu to compare one cell line with another cell line. Tumor subtype–specific loops were then merged using BEDTools pair to pair function with a negative slop of 10 kb. Hi-C breakfinder pipeline ^58^ was used to identify large structural translocations, deletions, and inversions. To identify neo-loops or enhancer hijacking events, we used a NeoLoopFinder pipeline as previously described algorithm ^35^.

### Targeting distal enhancers using CRISPRi and ChIP–qPCR

MB-004 cells were transduced with the enti-dCas9-KRAB-T2A-GFP virus (Addgene # 71237). sgRNAs targeting SOX11/HNRNPH1-binding motifs in the distal enhancers were designed using the CHOPCHOP program (https://chopchop.cbu.uib.no). GFP-reporter positive cells were flow-sorted 2 days transduction. DNA oligonucleotides were annealed and ligated into the lentiGuide-Cherry vector (Addgene # 170510) at the BsmBI restriction enzyme cutting sites. The sgRNA sequences were confirmed by Sanger sequencing. The lenti-vectors expressing each pair of gRNAs targeting distal enhancers were packaged in 293T cells using pMD2.G (Addgene # 12259) and psPAX (Addgene # 12260). The lentiGuide-Cherry lentiviruses carrying sgRNAs were concentrated from the virus-containing medium by ultracentrifuging and transduced into dCas9-KRAB-T2A-GFP-expressing MB-004 cells (MOI < 1). RNAs were then extracted from the GFP+/Cherry+ cells after 72 hr culture, and cDNAs were prepared using SuperScript^®^ III First-Strand Synthesis System (Invitrogen) according to the manufacturer’s instructions. qRT-PCR was performed to quantify gene expression using SYBR FAST qPCR Master Mix. All sgRNA and qRT-PCR sequences used for validation can be found in Supplementary Table 5.

ChIP-qPCR assays were performed as previously described ^59^. Briefly, cells (2 × 10^6^) were transduced with lentivirus expressing non-targeting shRNA control, shHNRHNP1, or shSOX11 for 60 hours (selected with puromycin) and then fixed with 1% formaldehyde for 10 min at room temperature and quenched with 0.2M glycine. Sonicated chromatin was prepared in buffer (10 mM Tris-HCl pH 8.0, 1 mM EDTA, 0.5 mM EGTA and protease inhibitor cocktail) and then incubated with 4 μg H3k27ac antibody (Abcam, ab4729) overnight at 4 C. Magnetic protein A/G beads were incubated to each ChIP reaction for 1 hr at 4 C. ChIP DNAs were eluted into 200 μl elution buffer at 65 °C for 20 min and extracted with phenol/chloroform. Purified DNAs were subjected to qRT-PCR assay for quantifying H3K27ac occupancy on the enhancers. Sequences of ChIP-qPCR primers are listed in Supplementary Table 5.

### Cut & Run-seq and data processing

Cut & Run-seq was performed as previously described^49^. Briefly, 200,000 cells were harvested and washed twice and captured by the addition of 10□μl pre-activated concanavalin A coated magnetic beads (Bangs Laboratories-BP531). Cells were then resuspended in 100 μl cold Antibody Buffer and 1□0μl antibody (H3K4me3, EpiCypher #13-0041; H3K27Ac, Active Motif #39133; SOX11, Sigma #HPA000536 or HNRNPH1 Abcam #ab154894) was added for incubating on nutator overnight at 4□°C. Cells were washed twice in 1□ml Digitonin Buffer (20 mM HEPES-KOH pH 7.5; 150□mM NaCl; 0.5□mM Spermidine; 1×Roche cOmpleteTM; 0.05% digitonin), and then resuspended with CUTANA pAG-MNase in 50□μl Digitonin Buffer. After wash twice, samples were quickly mixed with 100□mM CaCl_2_ to a final concentration of 2□mM and incubation at 4 °C for 30 min and reaction was quenched by Stop Buffer. The cleaved fragments were released by incubating the tube for 30□min at 37□°C and then purified by Qiagen MinElute PCR Purification Kit. Libraries were prepared with NEBNext^®^ Ultra™ II DNA Library Prep Kit for Illumina^®^ (E7645) and sequenced by NovaSeq PE150.

Cut & Run-seq reads were aligned to the reference human genome version hg38 with the program BOWTIE v2.3.4.1. Aligned reads were stripped of duplicate reads with the program sambamba v0.6.8. Peaks were called using the program MACS v2.1.2 with the narrow and broad peaks mode for Cut & Run-seq. Motif enrichment analysis was performed for both HNRNPH1- and SOX11-bound sites using HOMER findMotifsGenome function with -size 1000 -mask settings. TFs with high-expression level in G3 MB cell lines and significant H3K27Ac enrichment in G3 MB-specific enhancers were identified as G3 MB-specific active motifs.

### Statistics and reproducibility

All analyses in this research were performed using Microsoft Excel, GraphPad Prism 6.00 (San Diego California, https://www.graphpad.com) or RStudio (https://www.rstudio.com/ and R v.4.0.3, R Development Core Team, 2016). We use the “cor” function in R to calculate the Pearson correlation coefficient. Statistical significance was determined using two-tailed Student’s t tests as indicated. One-way ANOVA test was performed by multiple comparisons following Turkey’s ranking tests when comparing multiple groups. Data are shown as mean ± SEM. Values of p < 0.05 denoted a statistically significant difference. Quantifications were performed from at least three experimental groups in a blinded fashion. The n value was defined as the number of experiments that were repeated independently with similar results. No statistical methods were used to predetermine sample sizes, but our sample sizes are similar to those generally employed in the field. No randomization was used to collect all the data, but data were quantified with blinding.

Representative images/results of human fetal cerebella are shown from immunostaining on 2 independent fetal tissues with similar results at 9 PCW, 12 PCW, 14 PCW, and 16 PCW in Figure 2a-h and Extended data Figure 1f and 3a.

## Data availability

The high-throughput datasets generated in the current study are available in the Gene Expression Omnibus (GEO, GSE198565), European Genome-phenome Archive (EGA; https://www.ebi.ac.uk/ega/studies/) repositories (EGAD00001004435) and the raw data of single cell RNA-seq of human fetal cerebellum and medulloblastomas in the CNSA (https://db.cngb.org/cnsa/) of CNGBdb with accession number CNP0002781.

## Code availability

No custom code was used in this study. Open-source algorithms were used as detailed in analysis methods including Seurat 4.0.6 (https://satijalab.org/seurat/), Slingshot (https://bioconductor.org/packages/devel/bioc/vignettes/slingshot/inst/doc/vignette.html) Monocle 3 (https://cole-trapnell-lab.github.io/monocle3/), ArchR (https://www.archrproject.com/), CIBERSORTx (https://cibersortx.stanford.edu/), AltAnalyze (http://www.altanalyze.org/) and Vector (https://github.com/jumphone).

## Acknowledgements

The authors would like to thank Drs. Kathleen J. Millen and Ed Hurlock for comments, suggestions, and edits. We appreciate Arman Esshaghi Bayat for technical support. This study was funded in part by grants from the Cincinnati Research Foundation ARC award, CancerFree Kids foundation, Pray-Hope-Believe foundation, and TeamConnor Childhood Cancer Foundation to QRL.

## Contributions

Z.L., M.X., W.S., C. Z. J.W., and Q. R.L designed, performed and analyzed the majority of the experiments in this study. D.X., X.L., L.M.N.W. and S.C. contributed to the Cut&Run and partial immunostaining experiments. X.D., F.Z., K.B., R.R., R.X. and L.X. contributed to bioinformatics analysis and NMF workflow. Y.X., Y.L., W.M., S.T. and Q.X. contributed to primary tissue collection. S.O. contributed to sample collection and animal experiments. L.Z., X.W., F.Y., H.Z. and Y.L. contributed to HiC data analysis. M.X., C.B.S., P.D.B., J.P.P. and R.J.G. helped analyze data and provided inputs. J.M., H.L., W.Z., M.D.T and Q.R.L. provided the resources, guidance, and inputs. Z.L. and Q.R.L. wrote the manuscript.

## Extended Data Figure Legends

**Extended data Fig. 1 | Cell clustering analysis of human fetal cerebellar cells and NSC and TCP comparation. a**, Comparison of the number of genes/cell (left) and number of read counts/cell (right) between the sc-RNA-seq of human fetal cerebellar cells and published snRNA-seq dataset^5^. Unpaired two-tailed Student’s t test. **b,** UMAP clustering of human fetal cerebellum single cell samples. Colors indicate distinct populations from 95,542 single cells across eight developmental stages. **c**, *t*-SNE maps of cell populations at indicated developmental stages. One tissue at each stage was used for the scRNA-seq analysis. **d**, STREAM plot visualization of the cell density along the indicated cell lineage trajectories. **e**, Upper, a zoom-in view of lineage trajectories. Lower, volcano plot showing the differentially expressed genes between NSC and TCP. The genes were colored based on fold change differences (>1.5 folds) and FDR-p value (p < 0.05). The statistics was obtained by the two-sided Wilcoxon test as implemented by Seurat. **f**, Upper left, UMAP plots of NSC and TCP cell populations. Upper right and lower left panels indicate expression of G1/S and G2/M marker genes in NSC and TCP populations, respectively. Lower right, immunostaining for Ki67/HNRNPH1 in the RL transitional zone at PCW 12. Arrows: Ki67 and HNRNPH1 co-stained cells. Scale bar: 50 μm. **g,**Correlation analysis of the identified progenitor populations in the human fetal cerebellum.

**Extended data Fig. 2 | Cell type comparation of human fetal cerebella and mouse counterparts. a**, UMAP plot of single cells from human scRNA-seq (n = 95,542) and sn-RNA-seq (n = 69,174) from human fetal cerebella after LIGER analysis. The cells are colored by cell types. **b**, Riverplot comparing the human fetal cerebellar cell types from scRNA-seq with those from snRNA-seq dataset^5^ with the LIGER joint clustering assignments. **c**, UMAP plot showing only TCP (scRNA-seq), UBC_P (scRNA-seq) and RL (snRNA-seq) cells and colored by the cell types. **d**, Dot plot showing expression of NSC, TCP, UBC_P and RL marker genes from indicated human cerebellar scRNA-seq and snRNA-seq datasets. **e** UMAP plot of scRNA-seq data from human fetal cerebella (left) and mouse cerebella (right) after LIGER analysis. Cells are colored by cell types. **f**, Riverplot comparing the previously published mouse cerebellar cell clusters ^3^ and human fetal cerebellar cells identified from scRNA-seq with the LIGER joint clustering assignments. **g,** Dot plot showing expression of progenitor marker genes between human fetal and mouse cerebella.

**Extended data Fig. 3 | TCP and developmental hierarchy of cerebellar neural precursor cells. a**, The EGL region of PCW 12 fetal cerebellum stained for ATOH1 and HNRNPH1. Scale bars, 50 μm. Arrows indicate co-labeled cells. **b**, Upper, trajectory analyses from TCP to GCP (left), NSC to UBC (middle) and NSC to Purkinje lineages (right) in the human fetal cerebellum. Lower, Representative markers in the corresponding different populations. **c**, STREAM visualization of inferred branching points, trajectories and expression of key genes in the indicated cell populations.

**Extended data Fig. 4 | Trajectory analysis of human MB tumor cells. a**, CIBERSORTx deconvolution analyses of human fetal cerebellar populations against bulk transcriptomes of human MB subgroups from CBTTC cohort. **b-d**, Population, marker gene expression, and trajectory analysis of human MB tumors in G3 MBs (b, n = 52,743 cells), G4 MBs (c, n = 43,643 cells) or SHH MBs (d, n = 21,342 cells) using Vector based on the UMAP images. **e-h**, Reference map (e) of human fetal cerebellar neural progenitor cells based on the ProjecTILs profile. MYC^+^ G3 MB (f), G4 MB (g) and SHH MB(h) tumor cells were projected onto the reference map. **i**, Predicted fraction of cells in human MB subgroup CBTTC cohorts using CIBERSORTx deconvolution using SHH, G3 and G4 MB tumor cell type as reference.

**Extended data Fig. 5 | Distinct tumor cell populations in primary and metastatic medulloblastomas. a**, Scatter plots of mixed G3 and G4 MB tumor cells based on Group 3/4-B/C metaprogram expression scores. **b**, Heatmap of CNV signals normalized against the “Microglia” cluster shows CNV changes by chromosome (columns) for individual cells (rows) based on scRNA-seq data. **c-d**, UMAP of merged cells (c), split plot of cell populations (d), cell compositions (e) from the BT-325 primary and metastatic tumors. **f**, Two-dimensional cell-state plots and proportions of tumor subpopulations in BT-325 primary and metastatic MB. Each vertex corresponds to one key cellular state along the transformation. The colors represent the count. **g,** Dot plot displaying the expression of selected marker genes in the primary and metastatic tumors. The dot size reflects the percentage of the cells that express the gene. Average expression levels are color coded. **h**, CNV analysis of chromosome 8 based on DNA methylation showing the *MYC* gene amplification in the metastatic tumor of BT-325. **i**, Dot plot showing the expression levels of G3 and G4 marker genes in the BT-325 primary (x-axis) and metastatic (y-axis) tumors. Red and green indicate higher and lower expression, respectively, in the metastatic tumor compared to the primary tumor. **j**, UMAP of merged cells from the G3 MB (#2) primary and metastatic tumors (left). Split plot (middle and right) of cell populations from primary and metastatic samples. The cells are colored by cell type. **k**, Expression of indicated TCP markers and MYC in the cells from the G3 MB (#2) primary and metastatic tumors. **l, m**, Representative images (l) and the quantification (m) of four paired primary and metastatic human G3 MB tissues stained for HNRNPH1 and SOX11 (n = 4 individual patients). Scale bars: 50 μm. Unpaired two tailed t-test.

**Extended data Fig. 6 | Reference mapping of single-nuclei chromatin accessibility from G3 and G4 MBs. a**, Workflow for integrating scRNA-seq and snATAC-seq data analysis using ArchR. **b**, **c**, Comparation of the cell types in the b) G3 and c) G4 MB tumor tissues detected by scRNA-seq (upper) and snATAC-seq (lower) using Seurat and Signac analysis. **d**, **e**, The gene integration matrix from scRNA-seq and snATAC-seq (left panel) and alignment for key TCP- and MB-related marker genes (right panels) in d) G3 MB samples (n= 2 patients), and e) G4 MB samples (n = 3 patients). The individual cell types are indicated by colors. The integrated gene activity of G3 or G4 signature genes is shown in the right panels. **f**, Heatmap showing correlation of accessibility of promoter regions with target gene expression for tumor cell types. **g**, Pseudo-bulk ATAC-seq tracks at the indicated gene loci in G3 and G4 MB cells.

**Extended data Fig. 7 | Single-cell Omics reveals unique regulatory networks in transitional progenitor-like subpopulations in G3 and G4 MBs. a**, **b**, Heatmaps of scaled accessible peak-to-gene links identified along the integrated pseudotime trajectory from a) TCP-like to G3 MB, and b) TCP-like to G4 MB subclusters. Enriched transcription factor motifs are also shown. **c**, Dot plot showing positive transcriptional regulators identified in the TCP-like cells from G3 and G4 MB compared to other tumor subclusters. x-axis, the correlation of TF motif enrichment to TF gene expression. y-axis, ΔTF deviation scores between TCP-like cells and other subclusters. **d**, Tn5 bias-adjusted footprints for transcription factors SOX11 in TCP-like populations, MYC and OTX2 in G3 MBs, and LMX1A and RORA in G4 MBs. **e**, **f**, Genome accessibility track visualization of marker genes with peak co-accessibility in (e) TCP-G3-like cells in G3 MB (*OTX2* and *HLX*) and (f) TCP-like cells in G4 MB (*BARHL1* and *PAX6*).

**Extended data Fig. 8 | 3D genome organization differs between G3 and G4 MBs. a**, Signals of H3K27ac peaks at TSSs in G3-MB cells and NSC. **b**, Heatmap showing the relative density of indicated genes occupied by the H3K27ac, HNRNPH1, and SOX11 in G3 MB cells (MB-004) and human NSCs. **c**, Consensus motif analysis using HOMER for HNRNPH1 and SOX11 based on their genomic occupancy. The five highly represented motifs are shown. **d**, Heatmaps showing Hi-C interactions across genomes of G3 (left) and G4 (right) MBs. **e**, Aggregate peak analysis of G3-specific and G4-specific loops in G3 and G4 MB cells. **f**, **g**, the top-ranked structural variants identified by Hi-C mapping in G3 (f) and G4 (g) MB cells. **h**, Enhancer hijacking events identified in G3 MB-004 cells based on SV analysis. Inter-chromosomal translocation is observed between breakpoints chr11: 66,570,000 to 67,320,000 and chr19: 37,630,000 to 38,360,000. Genomic tracks are shown for H3K27ac/H3K4me3 occupancy in human NSCs and G3 MB cells. **i**, Expression of *PPP1R14A* in SHH, G3 and G4 MB cohorts based on Cavalli’s MB cohorts^31^. Data are shown as the mean ±□SEM (bars) and individual score (dots), two-tailed Student’s t test. **j**, Line graph presenting the number of superenhancers defined by ranked H3K27ac occupancy signal intensity using the ROSE algorithm defining 63 superenhancers in G3 MB-004 cells. **k**, Relative enrichment of H3K27ac occupancy on the enhancer or promoter of *OTX2* in MB-004 cells transduced with shCtrl, shSOX11 or shHNRNPH1, and in control hNSCs. n = 3 independent experiments for each region; unpaired two-tailed t-test.

**Extended data Fig. 9 | Targeting TCP signature genes alters the cell fate and inhibits tumor growth. a**, *HNRNPH11* (left) and *SOX11* (right) expression in subgroups of Cavalli’s cohort dataset. Data are shown as the mean ±□SEM (bar) and individual score (dots). **b**, Kaplan-Meier analysis of overall survival of patients with cerebellar MBs (WNT, SHH and G4 MBs) based on the TCP score from Cavalli’s MB cohorts. log-Rank test. **c**, Representative immunoblots for SOX11 and HNRNPH1 in human NSCs and G3 MB cell lines (D283, MB-004, and MB-002). n = 3 independent experiments. For gel source data, see supplementary Fig. 1. **d**, Heatmap showing gene expression profiles of control, shHNRNPH1 or shSOX11-transduced MB-004 cells. Representative signals or pathways, and the related genes are indicated. n = two independent experiments/treatment. Fold change > 1.5 folds and FDR *P*-value < 0.05. *P*-value was calculated by one-sided Fisher’s exact test adjusted with multiple comparisons. **e**, Heatmap showing that the genes associated with G3 MB signatures, TGF-beta signaling and EMT were significantly reduced by SOX11 or HNRNPH1 knockdown. **f,**Cell viability of indicated cells transduced with shHNRNPH1 (left) or shSOX11 (right) measured by WST-1 assay. n = 4 independent experiments. Data are shown as the mean ±□SEM. **g**, Representative immunostaining images of cleaved caspase 3 (upper) and quantification (lower) in MB-004 cells treated with shCtrl, shSOX11, or shHNRNPH1. n = 5 independent experiments per treatment. Data are shown as the mean ±□SEM (bars) and individual score (dots). Scale bars: 100 μm. **h**, Representative images of tumors in mice grafted with G3 MB-004 cells transduced with shCtrl, shSOX11, or shHNRNPH1. Representative tumors were collected (at day 18 post-implantation for Ctrl mice, at day 28 post-implantation for shSOX11 and at day 35 post-implantation for shHNRNPH1) and staining with hematoxylin and eosin (H/E) (left), Ki67 (middle) and cleaved caspase 3 (right). n = 5 independent samples/treatment. In a, f and g; two-tailed unpaired Student’s *t*-test.

**Extended data Fig. 10 | SOX11/HNRNPH1 expression in the mouse embryonic cerebellum and WNT MB. a**, **b**, Immunostaining images (a) and the quantification (b) for SOX11+/HNRNPH1+ cells in mouse embryonic cerebella at the indicated stage (n = 3 animals/stage). Scale bar: 200 μm. Data are shown as the mean ±□SEM (bars) and individual score (dots). VZ: ventricular zone, RL: rhombic lip. **c**, UMAP clustering of human WNT MB single cell RNA-seq dataset ^47^ Colors indicate distinct populations based on the gene expression profile. **d**, Expression of TCP signature genes (upper) and WNT MB-specific genes (lower) in WNT MB cells.

